# A Brain-wide Neuronal Spiking and Behavior Dataset for Working Memory-Specific Activation and Reactivation in Mice

**DOI:** 10.64898/2026.04.12.718071

**Authors:** Ermeng Huang, Da Xu, Huangao Zhu, Zhaoqin Chen, Peiyuan Li, Yulei Chen, Shujun Lei, Jiawei Liu, Chengyu Li, Xiaoxing Zhang

**Affiliations:** Institute of Neuroscience, State Key Laboratory of Neuroscience, CAS Center for Excellence in Brain Science and Intelligence Technology, Chinese Academy of Sciences, Shanghai 200031, China; Shanghai Center for Brain Science and Brain-Inspired Technology, Shanghai 200031, China; Lingang Laboratory, Shanghai, China

**Keywords:** Working Memory, Neural Networks, Neuroscience

## Abstract

How do millisecond-scale neural activities mediate multi-second behaviors such as working memory? This dataset provides brain-wide, high-spatiotemporal-resolution single-unit spiking activity recorded from 40 head-fixed mice performing an olfactory delayed paired-association (ODPA) task, encompassing 33,028 neurons across 62 anatomically defined brain regions. Utilizing this dataset enables the extraction of millisecond-scale within- and cross-region spike couplings enriched among memory-content-selective neurons. These couplings may be used to identify higher-order, hierarchically structured activity motifs. Because these multi-neuron patterns are captured both during active task performance and spontaneous inter-trial intervals, this dataset provides a resource for studying replay-related activity and for testing hypotheses about circuit-level processes associated with perceptual working memory. The dataset supports broad investigations into cross-regional spike organization, functional coupling, and temporal coding in working memory circuits.

## 1. Background and Summary

The brain operates across a spectrum of temporal scales, spanning from rapid millisecond-scale spiking activity to prolonged second-scale behaviors such as working memory ^[1–4]^. Working memory is a foundational cognitive function that bridges perception and action, with delay-period activity observed across various brain regions contributing to the active retention of task-relevant information ^[5–21]^. A central open question in systems neuroscience is how these millisecond-scale spike events are organized across regions to support such second-scale behaviors ^[18,21–31]^. Several theoretical models have been proposed to explain this, including the cell-assembly^[21–24]^ hypothesis of sequential activity in recurrent networks, the attractor network theory^[18,25–27]^ of sustained firing, the synfire chain model^[28–30]^ of feed-forward sequential spike propagation, and the concept of locally unstable yet globally stable population trajectories^[9,15,31,32]^. Furthermore, it is crucial to understand whether working memory representations are exclusively triggered by task demands or if they are spontaneously replayed beyond the boundaries of task execution, such as during inter-trial intervals ^[21,33–42]^.

This dataset provides brain-wide, high-spatiotemporal-resolution single-unit spiking data that can be used to investigate these questions. Using Neuropixels probes, electrophysiological recordings were collected from 40 head-fixed mice performing a delay-varying olfactory delayed paired-association working memory task. The dataset encompasses 33,028 neurons across 62 anatomically defined brain regions at the Allen Mouse Brain Common Coordinate Framework version 3 (CCFv3) level 7 ^[43]^, a level of granularity that provides a practical balance between group size and between-region specificity.

Serving as the dataset for our previously published findings on the hierarchical replay of multi-regional sequential spiking associated with working memory^[21]^, these recordings include both task-related neural activity and spontaneous recurrence of cross- regional activity patterns that can be analyzed in relation to prior findings. The dataset includes spike couplings and higher-order activity motifs that can be examined during the delay period and during inter-trial intervals. By making these extensive recordings publicly available, this dataset provides a resource for modeling replay-related dynamics, characterizing multi-second spiking activity organization, and evaluating hypotheses about circuit-level processes associated with perceptual working memory.

## 2. Data Records

### 2.1. Data title

A Brain-wide Neuronal Spiking and Behavior Dataset for Working Memory-Specific Activation and Reactivation in Mice

### 2.2 Data Content Description

116 data files containing spike times, quality control metrics, and brain region annotations for 33,028 spike-sorted single units, as well as behavioral event markers from a working memory task and preprocessed peri-stimulus time histograms (PSTHs; firing rates) aligned to the task structure.

### 2.3 Data Generation Time

October 2019 – January 2020

### 2.4 Data volume and data format

Approximately 17,323 MB (Sorted spike times, unit quality metrics, anatomy labels, and behavioral events; raw traces excluded), in the Neurodata Without Borders ‘.nwb’ format ^[44,45].^

### 2.5 Data source location

https://doi.org/10.57760/sciencedb.nb.00013

### 2.6 Citation format of data

Ermeng Huang, Da Xu, Huangao Zhu, et al. A Brain-wide Neuronal Spiking and Behavior Dataset for Working Memory-Specific Activation and Reactivation in Mice[DS/OL]. V1. Science Data Bank, 2026[2026-04-11]. https://doi.org/10.57760/sciencedb.nb.00013. DOI:10.57760/sciencedb.nb.00013.

### 2.7 Data Value

This dataset provides extensive brain-wide coverage, encompassing 33,028 single units across 62 brain regions, with 34 regions each containing more than 100 neurons. Its distinctive contribution lies in providing data suitable for analyzing both task-related activity motifs and their potential hierarchical reoccurrence during inter-trial intervals, a dimension that has been difficult to examine in prior working memory research. The dataset provides novel resources to:

- Characterize and constrain computational models of multi-second spiking activity organization underlying working memory, such as cell assemblies, attractor networks, synfire chains, and trajectory dynamics ^[18,22–31]^
- Investigate spike-coupled motifs and replay-related activity in relation to working memory, decision making, and long-term circuit-level plasticity ^[7,33–42,46–49]^
- Support the development of brain-inspired artificial intelligence architectures informed by hierarchical neural dynamics ^[32,50,51]^

The dataset is hosted on an open data platform for long-term preservation, adhering to established open-science best practices for data sharing.

### 2.8. Data Generation Method and Process

#### 2.8.1 Experimental Model and Subject Details

C57BL/6J Slac male adult mice were sourced from the Shanghai Laboratory Animal Center (SLAC), CAS, Shanghai, China. All subjects were male, aged 8–12 weeks, and weighed between 20–30 g at the initiation of the training protocols. Mice were socially housed in groups of 4–6 per cage within a tightly controlled environment under a 12-hour light-dark cycle (lights on from 6:30 a.m. to 6:30 p.m.). To ensure comprehensive brain- wide coverage, electrophysiological data were collected from 40 mice using varied probe trajectories; this broad sampling ultimately yielded more than 100 isolated neurons in each of 34 distinct brain regions. All experimental procedures strictly complied with standard animal care protocols and received approval from the Institutional Animal Care and Use Committee of the Institute of Neuroscience, Chinese Academy of Sciences (Shanghai, China, Reference Number NA-014-2019).

#### 2.8.2 Behavioral Setups

The behavioral paradigm utilized a custom-built olfactometry apparatus controlled by a programmable PIC Digital Signal Controller (dsPIC30F6010A, Microchip, Chandler, AZ). This controller precisely regulated olfactory cues and water delivery via solenoid valve switching and detected lick responses using a highly sensitive infra-red beam break detector. The 3D-printed odor delivery assembly comprised an air-and-odorant-mixture nozzle, a lick port, and the beam breaker assembly. This assembly was positioned directly in front of the head-fixed mice during all training and recording sessions (Fig. 1A).

**Figure 1.**
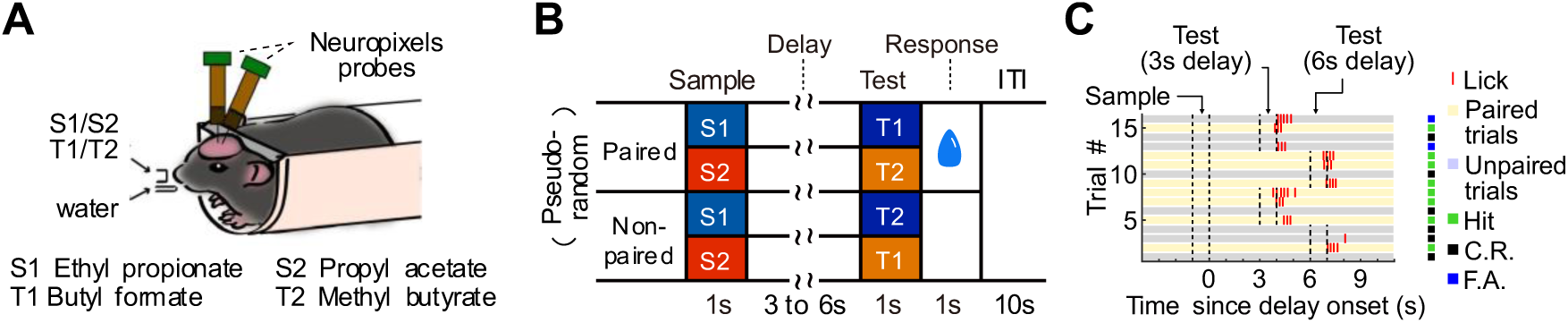
Behavioral and recording setup and task design. **A** Schematics of the head-fixed behavioral training and recording setup, detailing the specific chemical odorants utilized in the task. **B** Temporal structure of the olfactory delayed pair-association (ODPA) task, outlining the sample, delay, test, response window, and inter-trial interval (ITI) epochs. **C** Representative behavioral performance tracking over consecutive trials.

**Figure 2.**
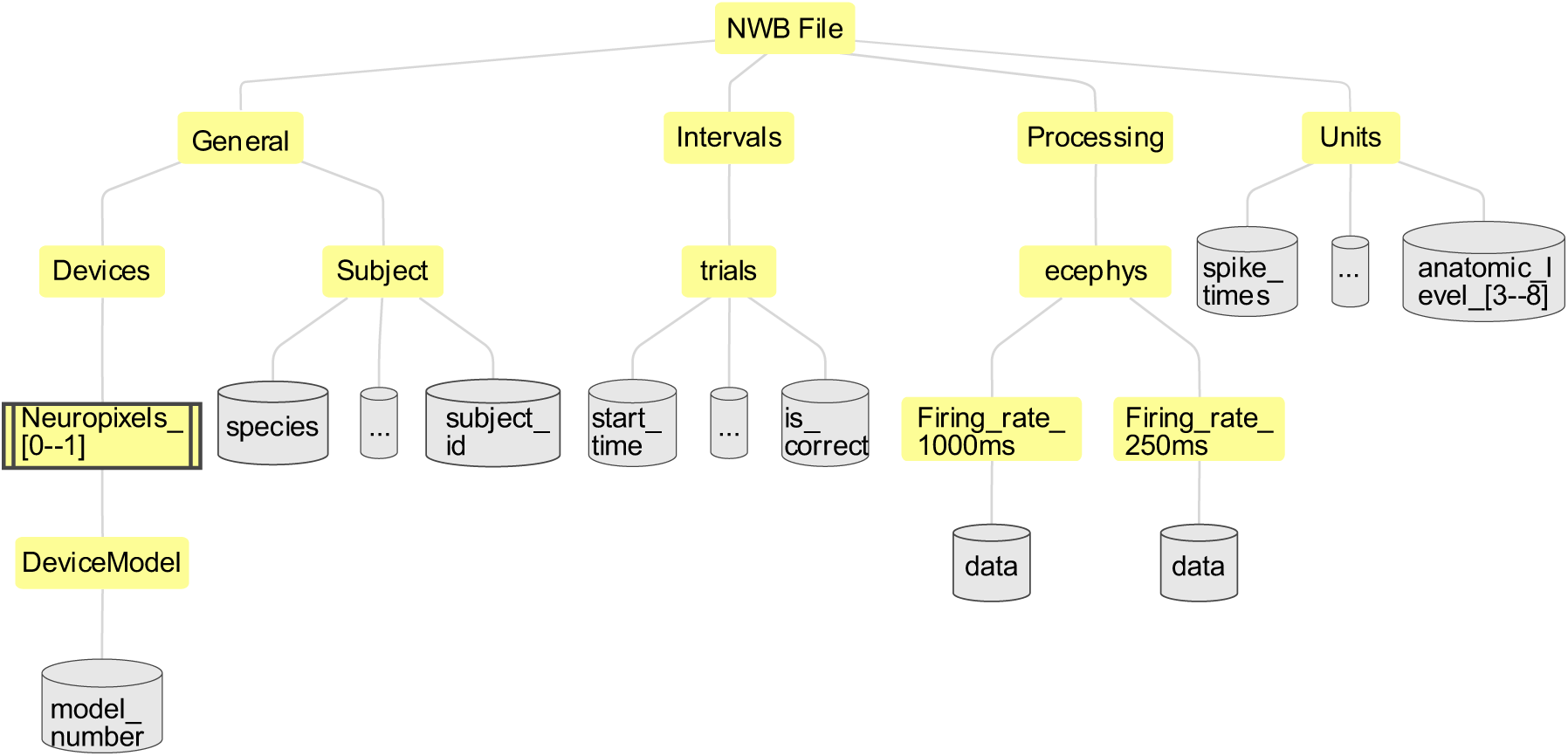
Hierarchical data structure of the Neurodata Without Borders (NWB) files. Each recording session is stored in an NWB file that is organized into four main data groups: **General:** Stores metadata for the subject (e.g., species, ID) and the recording devices, including specific Neuropixels probe models and hardware details. **Intervals:** Contains a trial-by-trial table specifying behavioral parameters, task event timestamps, and trial outcomes. **Processing:** Houses pre-computed trial-aligned neuronal firing rate time series binned at 1000ms and 250ms resolutions. **Units:** Stores neuronal data for sorted single units, encompassing spike times, cluster quality control metrics, and CCFv3 anatomical region annotations spanning parcellation granularity levels 3 through 8.

Air flow was maintained at a constant rate of 1.35 L/min. Liquid odorants including butyl formate (Cat. No. 261521), methyl butyrate (Cat. No. 246093), ethyl propionate (Cat. No. 112305), and propyl acetate (Cat. No. 133108; all from Sigma-Aldrich) were distributed in air-tight glass bottles to generate odorant vapor. Polytetrafluoroethylene (PTFE) tubes linked these bottles to the solenoid valves. During stimulus delivery, controlled air flow passed through the vapor bottles and was subsequently mixed with clean air at a 1:10 (v/v) dilution ratio. Odor concentrations were independently verified using a photoionization detector (200B miniPID, Aurora Scientific Inc.). Measurements confirmed that, within 1 second after valve shut-off, odor concentration returned to baseline levels, with residual concentration falling below 1% of the peak reached during cue delivery. Given that the working memory delay period lasted 3 s or 6 s, persistent sensory contamination during the delay was likely minimized.

#### 2.8.3 Behavioral Training Protocol

Mice performed the delay-varying olfactory delayed paired-association (ODPA) task (Fig. 1B)^[14,15,21]^. In this paradigm, one of two sample odors (S1: Ethyl propionate or S2: Propyl acetate) was presented for 1 second. This was followed by a variable delay period of either 3 seconds or 6 seconds, a design choice intended to reduce the association between working memory maintenance and reward expectation. Mini-blocks consisting of four consecutive trials alternated systematically between the 3-second and 6-second delay durations. Post-delay, one of two test odors (T1: Butyl formate or T2: Methyl butyrate) was presented for 1 second. Following the offset of the test odor, mice were granted a 1-second response window. The inter-trial interval (ITI) following the response window (and reward delivery when applicable) was set at 10 seconds.

The task required mice to learn specific pairings: S1 was paired with T1, and S2 was paired with T2. Mice were trained to lick the water port exclusively during the response window of paired trials. Behavioral outcomes were classified as follows: ‘hit’ and ‘false alarm’ were defined by licking during the response window of paired and unpaired trials, respectively. ‘miss’ and ‘correct rejection’ were defined by the absence of licking during paired and unpaired trials, respectively. Hits triggered an immediate 2.5 µL water reward, while false alarms, misses, and correct rejections yielded no reward or punishment. Daily training typically consisted of 60 blocks (240 trials total per session, Fig. 1C). Neural recordings were performed only after the mice had reached well-trained behavioral performance, defined as achieving at least 75% correct over 40 consecutive trials.

#### 2.8.4 Surgical Procedures and Neuropixels Implantation

Surgical preparation began with anesthetizing mice via intraperitoneal injection of sodium pentobarbital (10 mg/mL, 120 mg/kg body weight). Mice were secured in a stereotaxic frame equipped with a heating blanket. After removing the scalp and periosteum, a custom-designed stainless-steel head-plate was affixed to the posterior skull using tissue adhesive (1469 SB, 3M) and dental cement. Entry sites for the Neuropixels probes were marked, and a protective layer of dental cement was applied. Post-surgery recovery included consecutive 3-day intraperitoneal injections of antibiotics (0.2 mL 3.125 mg/mL ceftriaxone sodium) and a minimum 7-day recovery period before behavioral training commenced.

One day prior to electrophysiological recording, cranial windows (0.5–1.5 mm diameter) were drilled under isoflurane anesthesia without disrupting the dura mater. These windows were temporarily sealed with layers of 1.5% agarose, silicone elastomer, and dental cement. On the recording day, awake mice were head-fixed, the protective seals and dura mater were removed, and Neuropixels probes were inserted. To facilitate post hoc histological track reconstruction, each probe was systematically coated with fluorescent lipophilic dyes: DiI (red), DiO (yellow), and DiD (blue) (Invitrogen V22889, Thermo Fisher Scientific).

Probes were inserted to the predefined target depth and angle via the stereotaxic manipulator, typically at the speed of 1mm / minute while within brain tissue. A mandatory 30-minute stabilization period allowed the brain tissue to recover and minimized mechanical probe drift during the session. Baseline neural activity was recorded for 10 minutes prior to task initiation, and for an additional 10 minutes following task completion. Following extraction, probes were rigorously cleaned with filtered pure water (>18 MΩ·cm) and isopropyl alcohol.

#### 2.8.5 Electrophysiology Recording

The Neuropixels 1.0 (Order-code PRB_1_4_0480_1) is a fully integrated silicon CMOS extracellular recording probe^[52]^ designed for high-density electrophysiology in small laboratory animals, particularly rodents, where a single narrow shank can sample activity across multiple depths and often across several brain structures in one insertion. The probe combines on-shank amplification, filtering, multiplexing, and digitization, which reduces analog cabling requirements and supports stable acquisition of both spiking and low-frequency signals in standard neurophysiology workflows.

The device has one straight shank that is 10 mm long with a 70 × 24 µm cross-section, and carries 960 low-impedance TiN recording sites, of which only the lowest 384 were selected for simultaneous recording in this study. Each active channel enabled concurrent dual-band acquisition from the same selected sites. Data collection was primarily focused on the action-potential (AP) stream, which was sampled at 30 kHz with the band gain set to 1000. The local-field-potential (LFP) data were simultaneously acquired at 2.5 kHz with a band gain of 50, and were not used for further processing. The large TiN electrode located at the tip of the shank was used as reference electrode for each probe, across all sessions.

Data acquisition utilized SpikeGLX software (https://github.com/billkarsh/SpikeGLX) to simultaneously stream data from typically two Neuropixels probes. A custom-built embedded controller (based on a PIC12F1572 microcontroller, https://github.com/zhangxiaoxing/12F1572SyncEncoder) sampled task and behavioral signals at 1 MHz and generated a temporally encoded synchronization pattern of behavioral states at 500 Hz. This signal was routed directly into the SMA port of the Neuropixels base station, ensuring sub-millisecond alignment between neural events and behavioral states within the same auxiliary track of the binary file. Additionally, this signal provided a common reference time frame for high-precision synchronization across multiple probes.

#### 2.8.6 Spike Sorting

Kilosort 2 was used for fully automated spike sorting of the Neuropixels raw data^[53–55]^. Each recording was high-pass filtered at 150 Hz, and spikes were detected on each channel using a threshold of −6 times the noise standard deviation. The recordings were segmented into short temporal batches by the Kilosort 2 algorithm, which performs continuous drift correction and template tracking to follow putative single units across slow tissue movement along the probe shank. Spikes were assigned to clusters by template matching, and Kilosort 2 computed cluster-wise quality metrics, including contamination rate, firing rate, and template amplitude, for all putative units.

To standardize unit selection across sessions, we adopted Kilosort 2’s built-in, quantitatively defined criteria for labeling ‘good’ putative single units based on its internal quality metrics. Specifically, clusters had to satisfy a maximum contamination rate, exhibit a statistically significant refractory trough, and maintain a minimum mean firing rate over the entire recording. The exact thresholds and rationale are detailed in Section 2.10.2 (Neural Sorting Quality), together with the resulting distributions across units (Fig. 3B–D). By relying on Kilosort 2’ss automated metrics and fixed inclusion thresholds, we avoided manual cluster curation and ensured reproducible putative single-unit selection across all sessions.

**Figure 3.**
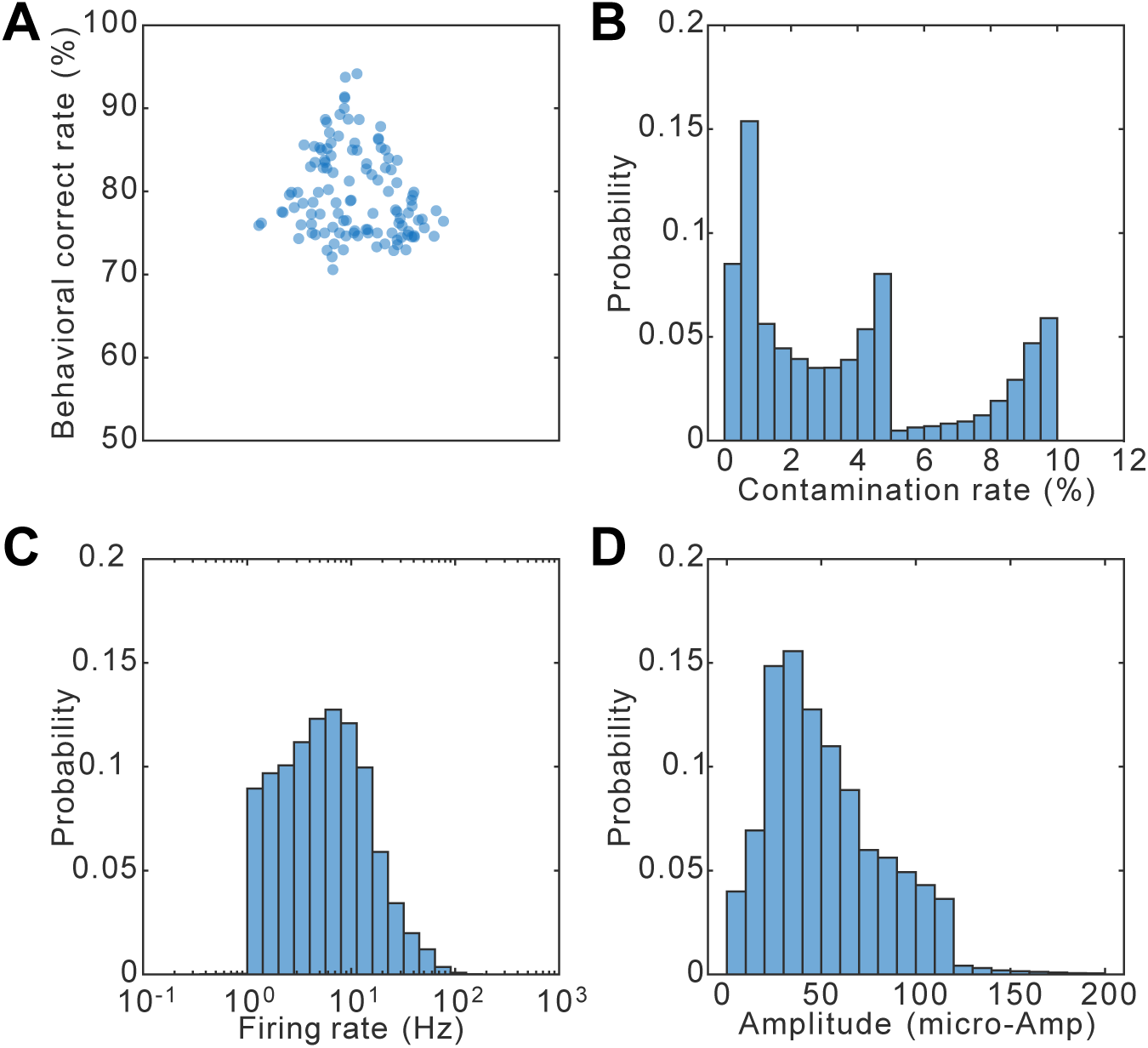
Behavioral and spike sorting data quality metrics. **A.** Behavioral performance (correct rates) of the mice in the task. **B-D** Spike sorting metrics for curated single units recorded in the behavioral task. **B.** Distribution of contamination rates. The discontinuity near 5% arises from the stepwise adaptive tightening of the contamination criterion in Kilosort 2 (see below). **C.** Distribution of mean firing rates for all included units. The sharp lower bound at 1 Hz arises from the minimum firing-rate threshold applied during putative single-unit selection. **D.** Amplitude of the mean spike waveform.

For sessions where two probes were recorded simultaneously, the sorted spike times from the second probe were linearly scaled and offset-adjusted to align perfectly with the time frame of the first probe, using the shared behavioral synchronization signal.

All Kilosort 2 outputs and derived metrics are stored in the NWB units table (see Section 2.9.5).

#### 2.8.7 Histology and Common Coordinate Framework Registration

Following terminal experiments, mice were deeply anesthetized and transcardially perfused with 0.9% NaCl followed by 4% paraformaldehyde (PFA) in phosphate-buffered saline (PBS). Brains were extracted, post-fixed overnight, and sectioned coronally into 80–100 µm slices using a vibratome. Sections were DAPI-stained and imaged at 10× magnification using an Olympus VS-120 fluorescence microscope across four channels (DiD: 645 nm; DiI: 535 nm; DiO: 488 nm; DAPI: 365 nm). The fluorescent lipophilic dyes (DiI, DiO, DiD) coated on the probe shanks produced clear tracks that could be traced across consecutive sections.

We employed a localization workflow modified from established protocols to reconstruct the anatomical trajectories of the recording probes^[56,57]^. Raw histological images were first downscaled to 20% of their original resolution. The brightness and contrast for each color channel were manually calibrated to maximize the visibility of both the fluorescent probe dye and the surrounding neuroanatomy. Images were then preprocessed and converted into TIFF format using a custom ImageJ macro (NPpreprocess.txt, included the pipeline below). Subsequent registration to the CCFv3 was performed using a custom open-source pipeline (https://github.com/StringXD/SmartTrack-Histology). Individual tissue slices were dimensionally scaled to align with the CCFv3 reference template, and approximately 10 paired anatomical landmarks were manually placed on both the histological images and the corresponding CCFv3 atlas sections to drive a non-linear warping procedure.

Because each probe insertion typically traversed multiple consecutive tissue slices, we manually annotated the visible fluorescent marks across all relevant sections and applied a linear best-fit approximation to model the continuous, straight-line probe trajectory in three dimensions. To integrate the neural data with this anatomical reconstruction, we matched each electrophysiological recording session to its corresponding physical track using experimental metadata, including subject ID, fluorescent dye color, stereotaxic insertion coordinates, and entry angle. The precise alignment of individual recording sites along each track was refined by (1) scaling the histological probe depth measurement—from the tip to the highest channel with biological signal—to match the physical tissue track depth, and (2) anchoring the distribution of electrophysiological landmarks (such as multi-unit activity transitions, single-unit spatial density peaks, and distinct PSTH patterns) across the evenly spaced recording sites to corresponding anatomical boundaries.

The finalized CCFv3-registered anatomical coordinates for each putative single unit—defined as its peak-amplitude recording channel—were exported as a standardized list and used to assign multi-level CCFv3 region labels. These labels were then embedded directly into the NWB units table to support downstream region-specific analyses (see Section 2.9.6). The overall confidence and limitations of this anatomical registration are further discussed in Section 2.10.3 (Histological Registration Quality).

To maintain high quality in our spatial registration, a subset of single units did not meet our criteria for accurate anatomical coordinate registration due to experimental limitations, such as a lack of clear fluorescence-labeled tracks or structural damage to the brain tissue during histological processing. These units were intentionally excluded from the regional mapping pipeline. Consequently, they do not possess specific CCFv3 region labels but are retained in the broader dataset for global statistical analyses.

### 2.9. Data sample description

All data associated with this release is available in Neurodata Without Borders (NWB) format ^[44,45]^. NWB has emerged as the premier standard for neurophysiology. It offers a robust ecosystem of open-source programming tools for efficient data access and analysis, and is widely adopted by major public repositories such as the Database of Brain Computer Interface at Lingang Laboratory, and the Distributed Archives for Neurophysiology Data Integration (DANDI), to streamline large-scale data sharing.

One NWB file is provided for each recording session. The filenames follow a BIDS-compatible naming convention of the form `sub-m<MOUSE_ID>_ses-`

`<DATE>_task-DPA.nwb` ^[58]^. The task ID indicates the behavioral paradigm performed (DPA: Delayed Paired-Association olfactory task). Each file contains an `identifier` field containing a string of this same form, where the mouse ID is a unique index into the internal data structures (e.g., m001, m002, m003) and the session date is encoded as an 8-digit number (YYYYMMDD).

#### 2.9.1 Structure of the NWB File

Each file contains the complete data from a single recording session for one subject. All files conform to the Neurodata Without Borders (NWB) core schema, version 2.9.0. These files can be accessed with standard NWB application programming interfaces available across major operating systems and programming environments, including PyNWB for Python and MatNWB for MATLAB. Because NWB is implemented on top of Hierarchical Data Format version 5 (HDF5), the files can also be read with general HDF5 libraries, provided that the NWB data organization and conventions are respected.

An NWB file comprises groups (like a directory) that store different subsets of data with pre-defined names and data types. For our dataset, we used the containers `general` (metadata for subjects and recording devices), `intervals` (task event timestamps and trial-specific behavioral flags), `processing` (pre-computed firing rate time series), and `units` (spike times of putative single units). The `description` field in each container was utilized to give further information on the structure and usage of the data. At the top level of each NWB file, additional metadata is stored in the fields `identifier`, `session_description`, `session_start_time`, and `timestamps_reference_time`, etc. All data are stored in strict accordance with the NWB standard and were validated using the NWB Inspector (https://nwb.org/tools/core/nwbinspector/) prior to public release.

#### 2.9.2 NWB File Content: General Group

The `general` group contains metadata. At the top level of `general`, metadata about the experiment is stored in the fields `experimenter`, `institution`, `lab`, `keywords`, `related_publications`, and `session_id`. Subject metadata is provided in the `general\subject` subgroup, including the subject’s species, strain, sex, age, genotype, and subject ID. Recording device information is provided in `general\devices`. A `DeviceModel` container specifies the probe manufacturer, model number, and description. For each individual Neuropixels probe recording data, a `Device` subgroup is created with a description, serial number, and a link to the DeviceModel.

#### 2.9.3 NWB File Content: Intervals Group

The `intervals` group specifies the trials in the session. Data is contained as a column-based table in the subgroup `intervals\trials` (accessible via `nwb.intervals.trials` in PyNWB or `nwb.intervals_trials` in MatNWB). There is one row for every trial. The `id` column references each trial (0-indexed).

For each trial, the following behavioral and task parameters are stored:

| Column | Description | Data Type | Notes |
| --- | --- | --- | --- |
| `start_time` | Start time of the trial | Float | In seconds relative to session onset |
| `stop_time` | Stop time of the delay period | Float | In seconds relative to session onset |
| `start_timestamp` | Start time of the trial in hardware sampling clock ticks | Integer | Clock ticks at 30000 Hz sampling rate |
| `stop_timestamp` | Stop time of the delay period in hardware sampling clock ticks | Integer | Same timebase as `start_timestamp` |
| sample_cue | Type of the sample cue odor presented | Integer | Value 4 represents S1 (Ethyl propionate); value 8 represents S2 (Propyl acetate). |
| test_cue | Type of the paired test cue odor | Integer | Value 8 represents T1 (Butyl formate); value 4 represents T2 (Methyl butyrate). |
| is_correct | Logical indicating if the trial outcome was correct | Logical | 1=correct, 0=incorrect |
| lick_response | Lick occurrence in the response window (or lack thereof) | Integer | 1=lick within response window, -1=no lick |
| delay_duration | Duration of the delay period | Integer | 3 or 6 seconds |
| well_trained | Indicator for trials within the well-trained behavioral window | Logical | 1=well-trained session, 0=disqualified session |

**Note:** In this specific task design, paired trials always consist of a sample cue value of 4 followed by a test cue value of 8, or vice versa. All NWB-standard time fields have been converted from raw hardware ticks to seconds to align with a unified session timebase, as required by the NWB schema. Original raw sampling ticks are retained in an auxiliary column for reference.

#### 2.9.4 NWB File Content: Processing Group

The `processing` group contains pre-computed analysis products. A `ProcessingModule` named `ecephys` contains time series data representing per-trial firing rates in specific time bins:

| TimeSeries | Description | Binning | Sampling Rate | Unit | Access Path |
| --- | --- | --- | --- | --- | --- |
| Firing_rate_1000ms | Per-trial firing rate with 1000ms bins, offset by -3 seconds | 1000ms | 1 Hz | Hz | nwb.processing.ecephys.Firing_rate_1000ms |
| Firing_rate_250ms | Per-trial firing rate with 250ms bins, offset by -3 seconds | 250ms | 4 Hz | Hz | nwb.processing.ecephys.Firing_rate_250ms |

These firing rate time series are aligned to trial events and can be used for rapid visualization and analysis without recomputing from raw spike times.

The alignment reference point for the “-3 seconds offset” is relative to sample cue onset (i.e. the first bin in each trial starts at 3 seconds before sample cue onset). Firing rates were computed for each unit and stored in a flattened matrix in accordance with NWB conventions. To maintain trial and temporal alignment, the first row records the trial index, the second row records the within-trial time-bin index, and each subsequent row contains the firing-rate series for an individual unit. The units are in the same order as in the units group (see below).

#### 2.9.5 NWB File Content: Units Group

The `units` group contains all the neuronal data (sorted single units). The data in `units` is stored in a column-based DynamicTable (accessible via `nwb.units`), with one row for each neuron (cluster). The `id` column denotes the per-session cluster index (0 based) and the `Cluster_id` denotes the cluster index as originally appeared in the Kilosort produced files for reference. Both indices are globally unique across multiple probe recordings within a session. Since Kilosort takes input of data from a single probe, when multiple probes are present within the same session, Cluster_ids from subsequent probes are offset by 10000 to ensure uniqueness.

Spike times are stored in `spike_times`, a region-based jagged array much larger than the number of rows in the units table, as it is a concatenation of all spike times across all cells. In accordance with NWB schema specifications, spike times are stored as floating-point numbers representing seconds relative to the start of the recording session, having been converted from raw 30,000 Hz hardware sampling ticks. The `spike_times_index` column contains indices to `spike_times`; each entry indexes the last spike of each neuron. Using this referencing scheme, the range of spikes for each neuron can be determined by using the value in `spike_times_index` as the ending value and the previous value in `spike_times_index` plus 1 as the starting index. Using `MatNWB`, the `nwb.units.toTable()` function can be used to conveniently extract all spike times—along with their corresponding unit IDs, quality control parameters, and anatomical localizations—into a single, unified MATLAB table.

#### 2.9.6 Quality Control Parameters

Additional metrics related to the quality of these clusters and their electrophysiological properties are provided:

| Column | Description | Data Type | Notes |
| --- | --- | --- | --- |
| `Cluster_id` | Globally unique cluster identifier | Integer | Offset by 10000 for the 2nd probe in multi-probe sessions |
| `KS_Label` | Kilosort2 quality classification | String | Possible values: "good", "mua", assigned by Kilosort. Only 'good' units are included in the current dataset |
| `Amplitude` | Cluster amplitude metric | Float | Spike amplitude in the physical signal space, unit: uV |
| ContamPct | Contamination percentage estimating spike sorting quality | Float | Percentage of contamination spikes in the refractory period |
| amp | Template amplitude from Kilosort2 | Float | Cluster template amplitude during sorting, not linearly related to spike amplitude, for reference only |
| channel | Channel number on the Neuropixels probe | Integer | 0-383 for NP1.0 |
| depth | Depth of the electrode site along the probe | Float | Distance in $\mu\text{m}$, from the probe tip to the site of maximum signal of a cluster |
| firing_rate | Mean firing rate in spikes per second | Float | Calculated as number of spikes divided by session duration |
| n_spikes | Total spike count for the cluster | Integer | Across entire session |
| shank | Shank number (for multi-shank probes) | Integer | 0 for NP1.0 (single shank) |

#### 2.9.7 Anatomical Localization

Brain region annotations are embedded directly in the units table as CCFv3 region labels at levels 3 to 8 of granularity (see Section 2.8.7 for the histological registration pipeline and Section 2.10.3 for a discussion of anatomical registration quality) ^[43]^. Levels 1 and 2 of the CCFv3 hierarchy are always “root” (abstract top-level container) and “grey” (grey matter) for all units and are therefore not stored explicitly. For each putative single unit, the anatomical hierarchy is represented by a set of string fields giving the CCFv3 acronym at each level:

| Column | Description | Data Type | Notes |
| --- | --- | --- | --- |
| `anatomic_level_3` | CCFv3 brain region at level 3 | String | e.g., "CH" (Cerebrum) |
| `anatomic_level_4` | CCFv3 brain region at level 4 | String | e.g., "CTX" (Cerebral cortex) |
| `anatomic_level_5` | CCFv3 brain region at level 5 | String | e.g., "CTXpl" (Cortical plate) |
| `anatomic_level_6` | CCFv3 brain region at level 6 | String | e.g., "HPF" (Hippocampal formation) |
| `anatomic_level_7` | CCFv3 brain region at level 7 | String | e.g., "HIP" (Hippocampal Region) |
| `anatomic_level_8` | CCFv3 brain region at level 8 | String | e.g., "DG" (Dentate gyrus) |

These annotations enable researchers to rapidly filter neural populations by specific structural compartments or to aggregate across levels of the Allen hierarchy without reprocessing histological images. Users who wish to adopt more conservative anatomical groupings can, for example, restrict analyses to higher-level labels (e.g., levels 3–5) or to regions with sufficient neuron counts, as summarized in the regional distribution table below.

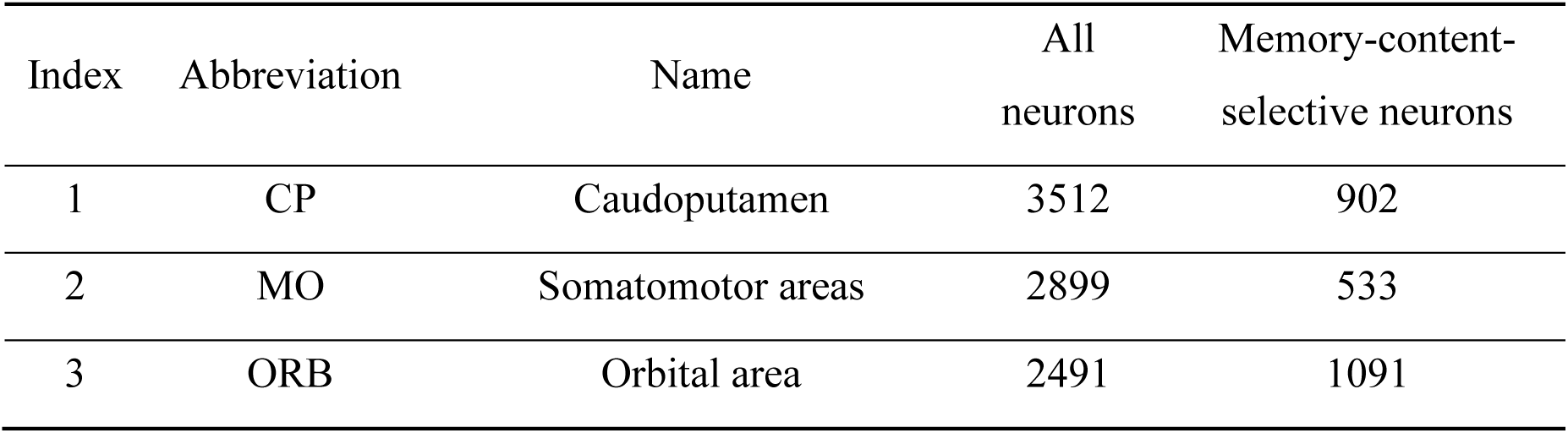
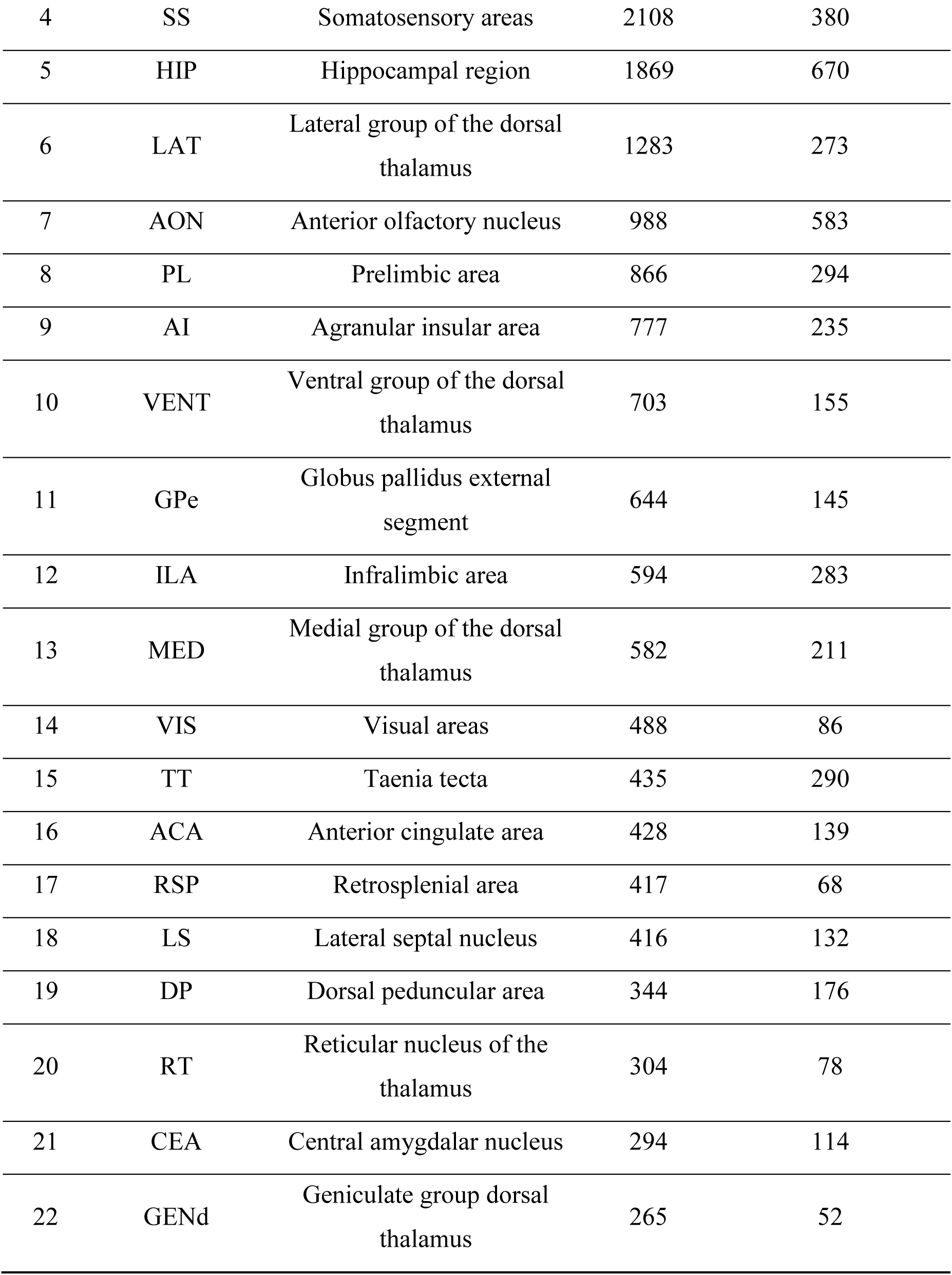
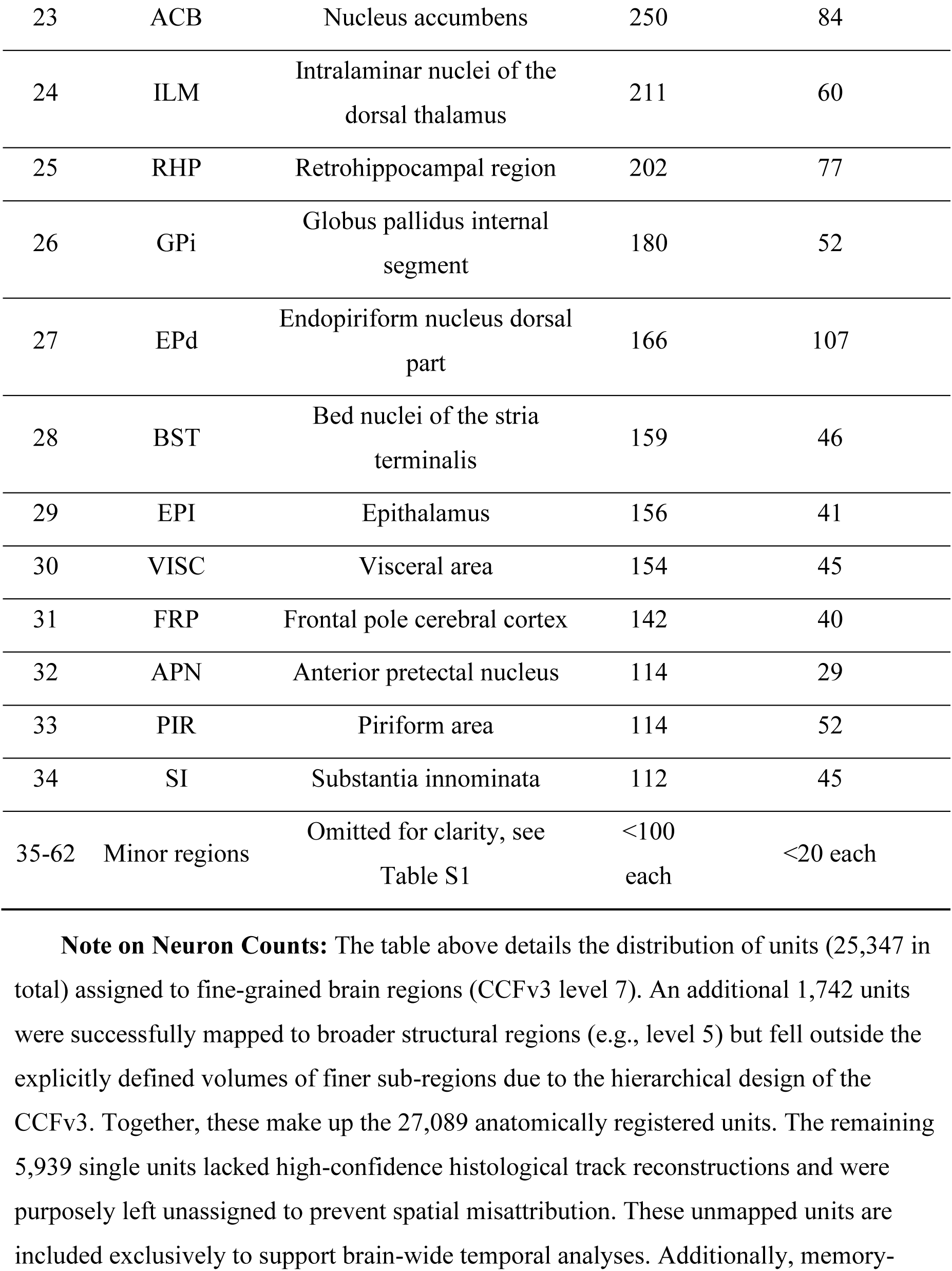
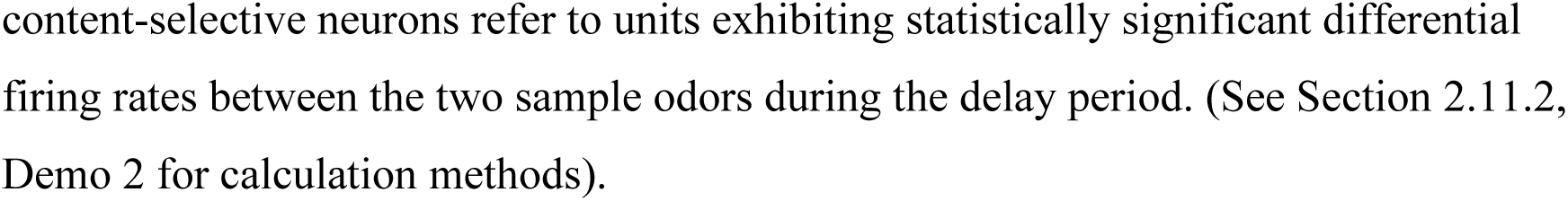

#### Note on Neuron Counts

The table above details the distribution of units (25,347 in total) assigned to fine-grained brain regions (CCFv3 level 7). An additional 1,742 units were successfully mapped to broader structural regions (e.g., level 5) but fell outside the explicitly defined volumes of finer sub-regions due to the hierarchical design of the CCFv3. Together, these make up the 27,089 anatomically registered units. The remaining 5,939 single units lacked high-confidence histological track reconstructions and were purposely left unassigned to prevent spatial misattribution. These unmapped units are included exclusively to support brain-wide temporal analyses. Additionally, memory-content-selective neurons refer to units exhibiting statistically significant differential firing rates between the two sample odors during the delay period. (See Section 2.11.2, Demo 2 for calculation methods).

### 2.10. Data Quality

#### 2.10.1. Behavioral Inclusion Criteria

The criterion for well-trained performance was defined as an accuracy of at least 75% over a moving window of 40 consecutive trials, consistent with previous studies using similar behavioral paradigms^[11,14,59,60]^. Neural-recording experiments were conducted only after mice reached this well-trained criterion; accordingly, the dataset includes only sessions obtained from well-trained mice. The mean behavioral accuracy within these well-trained windows is illustrated in Fig. 3A. Note that depending on the exact temporal distribution of correct and error trials, the overall mean accuracy may fall slightly below the moving-window threshold.

For future data reuse, we recommend that analyses of correct behavior are restricted to hit and correct-rejection trials occurring within the well-trained windows. Conversely, analyses of incorrect behavior may incorporate error trials from the entire recording session, encompassing periods both within and outside these well-trained windows.

#### 2.10.2. Neural sorting quality

As described in Section 2.8.6 (Spike Sorting), all recordings were processed with Kilosort 2 and cluster-wise quality metrics were computed automatically. Kilosort 2’s built-in, mathematically rigorous inclusion criteria—based on contamination rate, refractory-period ISI violations, and mean firing rate—were used to standardize ‘good’ unit selection across all sessions. Inter-spike interval (ISI) violations were quantified using an autocorrelogram (ACG) analysis: the central refractory trough (0–10 ms) was compared against expected baseline firing rates derived from the ACG shoulder regions (up to ±250 ms) under a Poisson null hypothesis. Clusters were classified as “good” putative single units only if the trough depletion was statistically significant (p ≤ 0.05) and the overall estimated contamination rate was ≤ 10%.

To prevent low-amplitude noise from degrading unit quality, Kilosort 2 iteratively evaluated contamination at progressively lower spike amplitude thresholds. If a putative unit demonstrated strong isolation (contamination < 5%) at the initial highest amplitude threshold, the algorithm adaptively tightened the maximum allowable contamination to ≤ 5% for all subsequent iterations, rejecting any lower amplitude threshold that would increase contamination above this stricter bound. All candidate clusters were additionally required to have a mean firing rate of ≥ 1 Hz across the entire recording session to ensure sufficient spike counts for robust template matching and reliable estimation of quality metrics.

These standardized parameters, rather than subjective human judgment, define the final set of putative single units in the dataset. Comparable fully automated approaches have also been adopted by other experienced research groups in the field ^[54,55]^. The resulting distributions of contamination rate, firing rate, and spike amplitude are summarized in Fig. 3B–D, demonstrating that the included units exhibit low estimated contamination and stable firing properties. Because all underlying quality metrics are stored in the NWB units table, downstream users can further tighten or relax these thresholds to match their analysis requirements or to benchmark against alternative spike-sorting pipelines.

#### 2.10.3. Histological registration quality

As described in Section 2.8.7 (Histology and Common Coordinate Framework Registration), each brain was registered to the CCFv3 using approximately 10 manually placed landmark pairs per slice and a *non-linear warping* procedure. This rich landmarking, combined with explicit tracing of fluorescent probe tracks across consecutive sections and linear best-fit modeling of the shank trajectory, provides high-confidence three-dimensional voxel placement for individual recording channels. Residual uncertainty is mainly expected near tissue boundaries, in regions with weaker fluorescent signal, or where local tissue deformation cannot be fully captured by the warping model.

To guarantee the reliability of spatial analyses, any single units originating from recording tracks that could not be unambiguously reconstructed were intentionally excluded from region-related analysis. This exclusion prevents spatial misattribution and ensures that downstream region-specific investigations are based on high-confidence anatomical ground truth. At the same time, preserving these unmapped units within the global NWB structures prevents the unnecessary loss of high-quality electrophysiological data that remains valuable for brain-wide temporal dynamics.

To accommodate different levels of anatomical precision required by downstream analyses, the dataset provides CCFv3 labels at multiple hierarchical levels (3–8) for each putative single unit (Section 2.9.6). Users who require conservative regional assignments may restrict analyses to higher-level CCFv3 labels (e.g., levels 3–5), or to regions with ample neuron counts (see the Distribution of Recorded Neurons Across Brain Regions table).

### 2.11. Data Usage Methods and Suggestions

#### 2.11.1. Recommended Software Environment

The dataset is provided in the Neurodata Without Borders (NWB) core schema, version 2.9.0. Because NWB is widely supported across programming languages, the data can be accessed using standard interfaces in both MATLAB and Python environments.

The analytical demonstration scripts accompanying this manuscript (Section 2.11.2) were developed in MATLAB, reflecting the primary analysis environment used in the original research study. To execute the provided scripts, we recommend the following setup:

#### MATLAB environment (required for provided scripts)

- MATLAB (Tested on R2025a).
- MatNWB – MATLAB interface for NWB file I/O (https://github.com/NeurodataWithoutBorders/matnwb/).
- FieldTrip Toolbox – For trial-aligned behavioral events and neural activity analysis (https://www.fieldtriptoolbox.org/).
- Buzsaki Lab Buzcode – For spike coupling analysis and cross-correlogram computation (https://buzsakilab.com/wp/resources/buzcode/).

While the current repository does not include Python equivalents of the analytical scripts, users working in Python can easily load and analyze the exact same NWB files using the standard ecosystem. A typical setup includes:

#### Python environment (for data access and custom analysis)

- Python 3.12 or later
- PyNWB – Python interface for NWB file I/O
- NumPy, pandas and SciPy – For numerical operations, metadata queries, and statistical analysis

All demo scripts provided with this dataset utilize MatNWB for file access. The NWB file structure enables direct access to spike times, trial tables, and anatomical annotations without requiring manual data parsing.

#### 2.11.2. Quick Start: Demonstration Scripts

To facilitate rapid adoption of this dataset, we provide three demonstration scripts illustrating common analysis workflows. The complete repository is openly available at https://gitee.com/XiaoxingZhang/hierarchical_replay_NSB_2025. These scripts serve as templates that can be adapted for custom analyses:

### Demo 1: Single-unit raster plot and peri-stimulus time histogram (PSTH)

- **Script:** jpsth/+demo/selective_unit_raster.m
- **Purpose:** Demonstrates how to load an NWB file, extract spike times for a single neuron, align spikes to task events, and generate raster plots with trial structure overlay.
- **Key operations:** NWB file reading via `nwbRead`, spike time extraction from `nwb.units`, trials table access from `nwb.intervals_trials`, pre-computed PSTH access from `nwb.ecephys`.
- **Expected output:** Figure showing spike rasters and PSTH aligned to sample cue onset for multiple trials (Figure 4A–B).

### Demo 2: Memory-selective neuron identification and regional distribution

- **Script:** jpsth/+demo/selective_ratio_per_region.m
- **Purpose:** Demonstrates statistical identification of memory-content-selective neurons (those exhibiting differential firing rates between sample odors during the delay period) and visualization of their distribution across brain regions.
- **Key operations:** Iterating over units, computing memory-selective responses, per-region selective-neuron proportion estimation with confidence intervals, CCFv3 region annotation access, and cross-region comparison.
- **Expected output:** Bar plot showing the proportion of memory-selective neurons per brain region (Figure 4C).

### Demo 3: Spike coupling detection and cross-correlogram analysis

- **Script:** jpsth/+demo/spike_coupling.m
- **Purpose:** Demonstrates identification of significant spike couplings between neuron pairs and computation of cross-correlograms (CCGs) using the baseline-corrected cross-correlation (BCCC) method adapted from previous publications ^[61–63].^
- **Key operations:** Simultaneous spike time extraction for selected neurons, trial selection based on behavioral criteria, coupled spike pair (CSP) detection within a defined latency window (<10 ms), raster visualization of coupled events, and CCG computation using the Buzcode toolbox.
- **Expected output:** Trial raster showing coupled spike events and cross-correlogram revealing asymmetric spike coupling (Figure 4D–E).

All demonstration scripts can be executed by invoking them in the working directory with the corresponding NWB data file. Users should first start MATLAB and add the MatNWB toolbox to the MATLAB path before running any demo.

**Figure 4.**
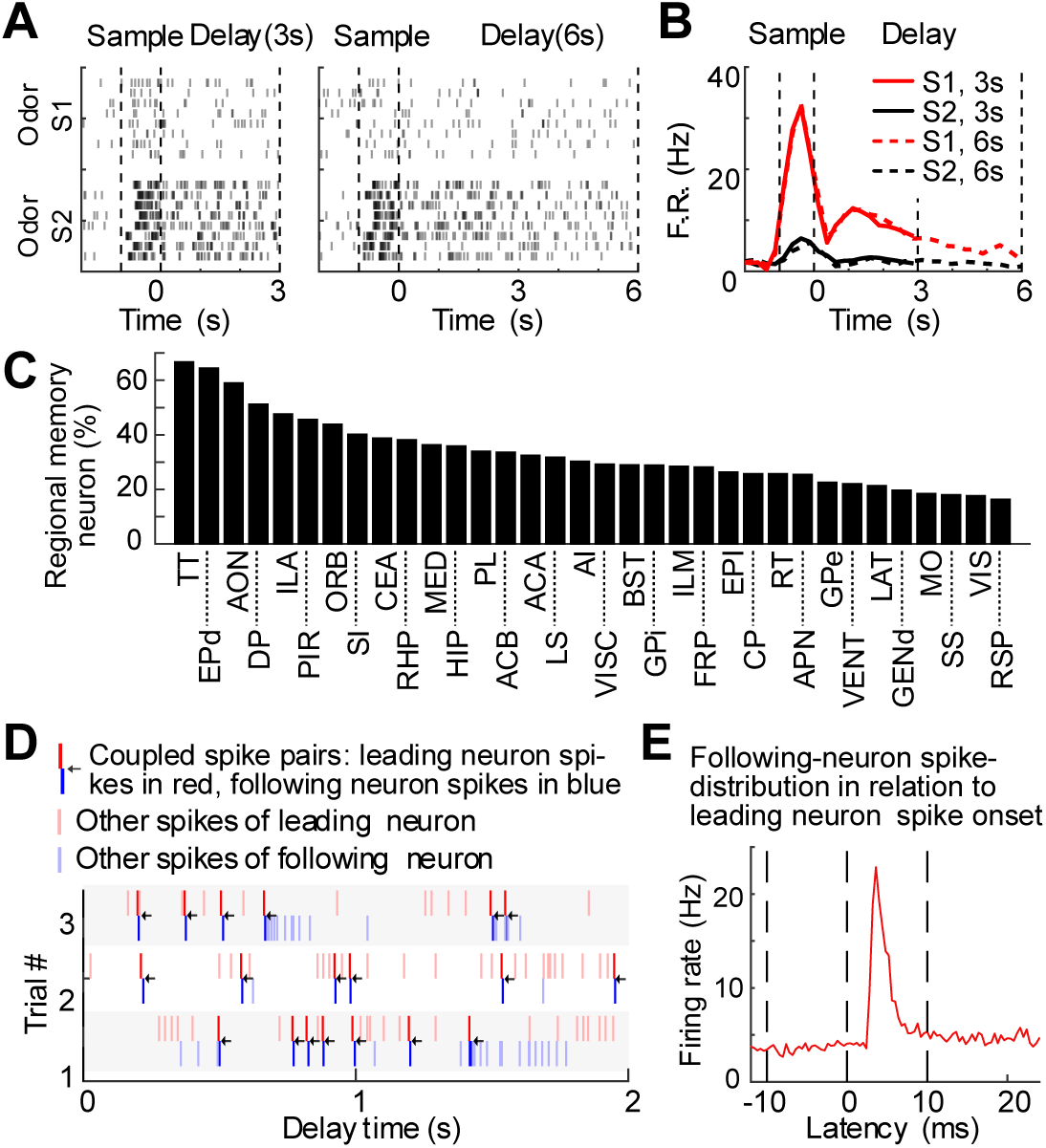
Data Usage Demonstration. **A** Activity of an example memory-content selective neuron (defined by the neurons with statistically significant different firing between two sample odors during the delay period). Left: Spike-raster plots of 8 example trials for two sample odors under 3-sec delay duration. Right: Spike raster for 6-sec delay duration. **B** Peristimulus time histogram (PSTH) of the example neuron in A. **C** Distribution of the proportion of memory-content selective neurons across 34 brain regions (only regions with more than 100 neurons registered are shown). **D** An example pair of simultaneously recorded neurons with spike coupling. Spike raster of three example trials from the two neurons with spike coupling (red for the leading neuron; blue for the following neuron). Arrows indicate the coupled spike pairs (CSP, with <10 msec latency). **E** Spike-correlogram showing the averaged spike frequency of the following neuron in relation to spiking of the leading neuron. Vertical dashed lines indicate the period used to determine asymmetric peaks in spike-correlogram.

#### 2.11.3. Data Access Best Practices

##### Trial selection

Behavioral trial outcomes are encoded in the `intervals_trials` table. For analyses of correct task performance, we recommend restricting to trials where both `well_trained` and `is_correct` values are `true`. For error analyses, trials from the entire session may be included. The `delay_duration` column distinguishes 3-second and 6-second delay conditions.

##### Temporal alignment

As required by the NWB core schema, all primary spike_times and trial start_time / stop_time fields are pre-converted to seconds. The original 30,000 Hz sampling clock ticks are preserved in auxiliary columns (e.g., start_timestamp, stop_timestamp). The sample cue onset serves as the primary behavioral alignment reference (time = 0) for peri-event analyses.

##### Anatomical queries

Brain region annotations are provided at six hierarchical levels of the CCFv3 parcellation (levels 3–8). Users can filter neurons by region using string matching on columns `anatomic_level_3` through `anatomic_level_8`. The Distribution of Recorded Neurons Across Brain Regions table provides the distribution of recorded neurons across regions for reference.

For additional support, users are encouraged to consult the NWB documentation (https://nwb.org) or report issues via the repository’s issue tracker.

## 3. Brief Data Overview

This dataset provides brain-wide single-unit spiking and behavioral data from 40 mice performing an olfactory working memory task, capturing 33,028 neurons across 62 brain regions through 116 sessions and 223 Neuropixels probe insertions. It includes cross-regional activity patterns, ranging from spike coupling to higher-order structured motifs, that can be analyzed during the delay period and inter-trial intervals. As a comprehensive open-access resource, the dataset enables investigation of millisecond-scale spiking activity associated with second-scale working memory behavior and provides an empirical foundation for computational modeling, neural circuit analysis, and brain-inspired algorithm development.

## 4. Data Availability Statement

This dataset is openly available via the Brain-Computer Interface database at Lingang Laboratory (https://dbci.lglab.ac.cn/) and ScienceDB platform (https://www.scidb.cn).

## 5. Code availability statement

The custom code used for analysis is available at https://gitee.com/XiaoxingZhang/hierarchical_replay_NSB_2025. The code was developed and tested in MATLAB R2025A. Required packages and tested versions are listed in the repository README.md.

## 6. Acknowledgments

We thank Drs. Muming Poo, Liping Wang, Bin Min, Yunzhe Liu, Ninglong Xu, Tianming Yang, Min Xu, Chun Xu, and Yong Gu for their critical comments on the project. We also thank the team responsible for the Brain-Computer Interface database at Lingang Laboratory, Dr. Jingpeng Wu, Chengqi Ke, and Yiqin Zhang for support with data-processing tools and data hosting. We further acknowledge the Center for Data and Computing in Brain Science, as well as the light microscopy and electron microscopy core facilities of CEBSIT/ION, for their support in generating the corresponding results. Jincan Hou managed the experimental animals used in this study. This work was supported by the Innovations of Science and Technology 2030 from the Ministry of Science and Technology of China 2021ZD0203601, 2022ZD0210300; National Natural Science Foundation of China 32221003 and 32161133024; Shanghai Municipal Science and Technology Major Project 2018SHZDZX05 and 2021SHZDZX; Shanghai Pilot Program for Basic Research–Chinese Academy of Sciences, Shanghai Branch JCYJ-SHFY-2022-010; National Key R&D Program Key Scientific Issues of Transformational Technology 2019YFA0709504; Lingang Laboratory Grant LGL-2930, LGL-1630, LG202105.

## 7. Author contributions

E.H., D.X., and H.Z. performed the behavioral and recording experiments, analyzed the data, and prepared the figures. Z.C., Y.C., S.L., and J.L. performed recording and additional experiments. P.L. assisted with the data-processing pipeline. X.Z. and C.L. conceived the project, supervised the experiments, analyzed the data, and wrote the manuscript.

## 9. Ethics statement

To achieve comprehensive brain-wide coverage, data were collected from 40 mice. All animal procedures were conducted in accordance with standard institutional guidelines for animal care and were approved by the Institutional Animal Care and Use Committee of the Institute of Neuroscience, Chinese Academy of Sciences (Shanghai, China, Reference Number NA-014-2019).

## References

[1] Goldman-Rakic P S. Cellular basis of working memory[J/OL]. Neuron, 1995, 14(3): 477–485. DOI:10.1016/0896-6273(95)90304-6.

[2] Fuster J M. The Prefrontal Cortex (Fifth Edition)[M/OL]. San Diego: Academic Press, 2015. DOI:10.1016/b978-0-12-373644-4.00008-6.

[3] Baddeley A. Working Memory, Thought, and Action[M/OL]. New York, NY, US: Oxford University Press, 2007. DOI:10.1093/acprof:oso/9780198528012.001.0001.

[4] Mckenzie S, Buzsáki G. Micro-, Meso- and Macro-Dynamics of the Brain[J/OL]. Research and Perspectives in Neurosciences, 2016: 1–21. DOI:10.1007/978-3-319-28802-4_1.

[5] Fuster J M, Alexander G E. Neuron Activity Related to Short-Term Memory[J/OL]. Science, 1971, 173(3997): 652–654. DOI:10.1126/science.173.3997.652.

[6] Baeg E H, Kim Y B, Huh K, et al. Dynamics of Population Code for Working Memory in the Prefrontal Cortex[J/OL]. Neuron, 2003, 40(1): 177–188. DOI:10.1016/s0896-6273(03)00597-x.

[7] Fujisawa S, Amarasingham A, Harrison M T, et al. Behavior-dependent short-term assembly dynamics in the medial prefrontal cortex[J/OL]. Nature Neuroscience, 2008, 11(7): 823–833. DOI:10.1038/nn.2134.

[8] Erlich J C, Bialek M, Brody C D. A Cortical Substrate for Memory-Guided Orienting in the Rat[J/OL]. Neuron, 2011, 72(2): 330–343. DOI:10.1016/j.neuron.2011.07.010.

[9] Harvey C D, Coen P, Tank D W. Choice-specific sequences in parietal cortex during a virtual-navigation decision task[J/OL]. Nature, 2012, 484(7392): 62–68. DOI:10.1038/nature10918.

[10] Guo Z V, Li N, Huber D, et al. Flow of Cortical Activity Underlying a Tactile Decision in Mice[J/OL]. Neuron, 2014, 81(1): 179–194. DOI:10.1016/j.neuron.2013.10.020.

[11] Liu D, Gu X, Zhu J, et al. Medial prefrontal activity during delay period contributes to learning of a working memory task[J/OL]. Science, 2014, 346(6208): 458–463. DOI:10.1126/science.1256573.

[12] Christophel T B, Klink P C, Spitzer B, et al. The Distributed Nature of Working Memory[J/OL]. Trends in Cognitive Sciences, 2017, 21(2): 111–124. DOI:10.1016/j.tics.2016.12.007.

[13] Pinto L, Rajan K, Depasquale B, et al. Task-Dependent Changes in the Large-Scale Dynamics and Necessity of Cortical Regions[J/OL]. Neuron, 2019, 104(4): 810–824.e9. DOI:10.1016/j.neuron.2019.08.025.

[14] Zhang X, Yan W, Wang W, et al. Active information maintenance in working memory by a sensory cortex[J/OL]. eLife, 2019, 8: e43191. DOI:10.7554/elife.43191.

[15] Zhu J, Cheng Q, Chen Y, et al. Transient Delay-Period Activity of Agranular Insular Cortex Controls Working Memory Maintenance in Learning Novel Tasks[J/OL]. Neuron, 2020, 105(5): 934–946.e5. DOI:10.1016/j.neuron.2019.12.008.

[16] Taxidis J, Pnevmatikakis E A, Dorian C C, et al. Differential Emergence and Stability of Sensory and Temporal Representations in Context-Specific Hippocampal Sequences[J/OL]. Neuron, 2020, 108(5): 984–998.e9. DOI:10.1016/j.neuron.2020.08.028.

[17] Wu Z, Litwin-Kumar A, Shamash P, et al. Context-Dependent Decision Making in a Premotor Circuit[J/OL]. Neuron, 2020, 106(2): 316–328.e6. DOI:10.1016/j.neuron.2020.01.034.

[18] Inagaki H K, Fontolan L, Romani S, et al. Discrete attractor dynamics underlies persistent activity in the frontal cortex[J/OL]. Nature, 2019, 566(7743): 212–217. DOI:10.1038/s41586-019-0919-7.

[19] Li N, Daie K, Svoboda K, et al. Robust neuronal dynamics in premotor cortex during motor planning[J/OL]. Nature, 2016, 532(7600): 459–464. DOI:10.1038/nature17643.

[20] Voitov I, Mrsic-Flogel T D. Cortical feedback loops bind distributed representations of working memory[J/OL]. Nature, 2022, 608(7922): 381–389. DOI:10.1038/s41586-022-05014-3.

[21] Huang E, Xu D, Zhu H, et al. Hierarchical Replay of Multi-regional Sequential Spiking Associated with Working Memory[J/OL]. Neuroscience Bulletin, 2025: 1–20. DOI:10.1007/s12264-025-01524-y.

[22] Hebb D O. The organization of behavior: A neuropsychological theory[M]. Psychology press, 2005.

[23] Harris K D. Neural signatures of cell assembly organization[J/OL]. Nature Reviews Neuroscience, 2005, 6(5): 399–407. DOI:10.1038/nrn1669.

[24] Buzsáki G. Neural Syntax: Cell Assemblies, Synapsembles, and Readers[J/OL]. Neuron, 2010, 68(3): 362–385. DOI:10.1016/j.neuron.2010.09.023.

[25] Hopfield J J. Neural networks and physical systems with emergent collective computational abilities.[J/OL]. Proceedings of the National Academy of Sciences, 1982, 79(8): 2554–2558. DOI:10.1073/pnas.79.8.2554.

[26] Zylberberg J, Strowbridge B W. Mechanisms of Persistent Activity in Cortical Circuits: Possible Neural Substrates for Working Memory[J/OL]. Annual Review of Neuroscience, 2017, 40(1): 603–627. DOI:10.1146/annurev-neuro-070815-014006.

[27] Wang X J. 50 years of mnemonic persistent activity: quo vadis?[J/OL]. Trends in Neurosciences, 2021, 44(11): 888–902. DOI:10.1016/j.tins.2021.09.001.

[28] Diesmann M, Gewaltig M O, Aertsen A. Stable propagation of synchronous spiking in cortical neural networks[J/OL]. Nature, 1999, 402(6761): 529–533. DOI:10.1038/990101.

[29] Ikegaya Y, Aaron G, Cossart R, et al. Synfire Chains and Cortical Songs: Temporal Modules of Cortical Activity[J/OL]. Science, 2004, 304(5670): 559–564. DOI:10.1126/science.1093173.

[30] Georgescauld D. Local Cortical Circuits, An Electrophysiological Study M. Abeles. Studies of Brain Function. Vol. 6. Springer-Verlag, Berlin, Heidelberg, New York, 1982, viii + 102 pp., DM39, US$18.20[J/OL]. Bioelectrochemistry and Bioenergetics, 1983, 10(2-3): 309. DOI:10.1016/0302-4598(83)85089-2.

[31] Rabinovich M, Huerta R, Laurent G. Transient Dynamics for Neural Processing[J/OL]. Science, 2008, 321(5885): 48–50. DOI:10.1126/science.1155564.

[32] Liu C, Jia S, Liu H, et al. Recurrent neural networks with transient trajectory explain working memory encoding mechanisms[J/OL]. Communications Biology, 2025, 8(1): 137. DOI:10.1038/s42003-024-07282-3.

[33] Liu Y, Nour M M, Schuck N W, et al. Decoding cognition from spontaneous neural activity[J/OL]. Nature Reviews Neuroscience, 2022, 23(4): 204–214. DOI:10.1038/s41583-022-00570-z.

[34] Kaefer K, Stella F, Mcnaughton B L, et al. Replay, the default mode network and the cascaded memory systems model[J/OL]. Nature Reviews Neuroscience, 2022, 23(10): 628–640. DOI:10.1038/s41583-022-00620-6.

[35] Swanson R A, Levenstein D, Mcclain K, et al. Variable specificity of memory trace reactivation during hippocampal sharp wave ripples[J/OL]. Current Opinion in Behavioral Sciences, 2020, 32: 126–135. DOI:10.1016/j.cobeha.2020.02.008.

[36] Skaggs W E, Mcnaughton B L. Replay of Neuronal Firing Sequences in Rat Hippocampus During Sleep Following Spatial Experience[J/OL]. Science, 1996, 271(5257): 1870–1873. DOI:10.1126/science.271.5257.1870.

[37] Wilson M A, Mcnaughton B L. Reactivation of Hippocampal Ensemble Memories During Sleep[J/OL]. Science, 1994, 265(5172): 676–679. DOI:10.1126/science.8036517.

[38] Nádasdy Z, Hirase H, Czurkó A, et al. Replay and Time Compression of Recurring Spike Sequences in the Hippocampus[J/OL]. The Journal of Neuroscience, 1999, 19(21): 9497–9507. DOI:10.1523/jneurosci.19-21-09497.1999.

[39] Foster D J, Wilson M A. Reverse replay of behavioural sequences in hippocampal place cells during the awake state[J/OL]. Nature, 2006, 440(7084): 680–683. DOI:10.1038/nature04587.

[40] Carr M F, Jadhav S P, Frank L M. Hippocampal replay in the awake state: a potential substrate for memory consolidation and retrieval[J/OL]. Nature Neuroscience, 2011, 14(2): 147–153. DOI:10.1038/nn.2732.

[41] Euston D R, Tatsuno M, Mcnaughton B L. Fast-Forward Playback of Recent Memory Sequences in Prefrontal Cortex During Sleep[J/OL]. Science, 2007, 318(5853): 1147–1150. DOI:10.1126/science.1148979.

[42] O’neill J, Boccara C N, Stella F, et al. Superficial layers of the medial entorhinal cortex replay independently of the hippocampus[J/OL]. Science, 2017, 355(6321): 184–188. DOI:10.1126/science.aag2787.

[43] Wang Q, Ding S L, Li Y, et al. The Allen Mouse Brain Common Coordinate Framework: A 3d Reference Atlas[J/OL]. Cell, 2020, 181(4): 936–953.e20. DOI:10.1016/j.cell.2020.04.007.

[44] Teeters J L, Godfrey K, Young R, et al. Neurodata Without Borders: Creating a Common Data Format for Neurophysiology[J/OL]. Neuron, 2015, 88(4): 629–634. DOI:10.1016/j.neuron.2015.10.025.

[45] Rübel O, Tritt A, Ly R, et al. The Neurodata Without Borders ecosystem for neurophysiological data science[J/OL]. eLife, 2022, 11: e78362. DOI:10.7554/elife.78362.

[46] Takeuchi D, Hirabayashi T, Tamura K, et al. Reversal of Interlaminar Signal Between Sensory and Memory Processing in Monkey Temporal Cortex[J/OL]. Science, 2011, 331(6023): 1443–1447. DOI:10.1126/science.1199967.

[47] Funahashi S, Inoue M. Neuronal Interactions Related to Working Memory Processes in the Primate Prefrontal Cortex Revealed by Cross-correlation Analysis[J/OL]. Cerebral Cortex, 2000, 10(6): 535–551. DOI:10.1093/cercor/10.6.535.

[48] Rao S G, Williams G V, Goldman-Rakic P S. Isodirectional Tuning of Adjacent Interneurons and Pyramidal Cells During Working Memory: Evidence for Microcolumnar Organization in PFC[J/OL]. Journal of Neurophysiology, 1999, 81(4): 1903–1916. DOI:10.1152/jn.1999.81.4.1903.

[49] Gupta A S, Meer M A A van der, Touretzky D S, et al. Hippocampal Replay Is Not a Simple Function of Experience[J/OL]. Neuron, 2010, 65(5): 695–705. DOI:10.1016/j.neuron.2010.01.034.

[50] Ven G M van de, Siegelmann H T, Tolias A S. Brain-inspired replay for continual learning with artificial neural networks[J/OL]. Nature Communications, 2020, 11(1): 4069. DOI:10.1038/s41467-020-17866-2.

[51] Wittkuhn L, Chien S, Hall-Mcmaster S, et al. Replay in minds and machines[J/OL]. Neuroscience & Biobehavioral Reviews, 2021, 129: 367-388. DOI:10.1016/j.neubiorev.2021.08.002.

[52] Jun J J, Steinmetz N A, Siegle J H, et al. Fully integrated silicon probes for high-density recording of neural activity[J/OL]. Nature, 2017, 551(7679): 232–236. DOI:10.1038/nature24636.

[53] Pachitariu M, Sridhar S, Pennington J, et al. Spike sorting with Kilosort4[J/OL]. Nature Methods, 2024, 21(5): 914–921. DOI:10.1038/s41592-024-02232-7.

[54] Stringer C, Pachitariu M, Steinmetz N, et al. Spontaneous behaviors drive multidimensional, brainwide activity[J/OL]. Science, 2019, 364(6437): 255. DOI:10.1126/science.aav7893.

[55] Luo T Z, Bondy A G, Gupta D, et al. An approach for long-term, multi-probe Neuropixels recordings in unrestrained rats[J/OL]. eLife, 2020, 9: e59716. DOI:10.7554/elife.59716.

[56] Steinmetz N A, Zatka-Haas P, Carandini M, et al. Distributed coding of choice, action and engagement across the mouse brain[J/OL]. Nature, 2019, 576(7786): 266–273. DOI:10.1038/s41586-019-1787-x.

[57] Shamash P, Carandini M, Harris K, et al. A tool for analyzing electrode tracks from slice histology[J/OL]. bioRxiv, 2018: 447995. DOI:10.1101/447995.

[58] Gorgolewski K J, Auer T, Calhoun V D, et al. The brain imaging data structure, a format for organizing and describing outputs of neuroimaging experiments[J/OL]. Scientific Data, 2016, 3(1): 160044. DOI:10.1038/sdata.2016.44.

[59] Han Z, Zhang X, Zhu J, et al. High-Throughput Automatic Training System for Odor-Based Learned Behaviors in Head-Fixed Mice[J/OL]. Frontiers in Neural Circuits, 2018, 12: 15. DOI:10.3389/fncir.2018.00015.

[60] Yao J, Hou R, Fan H, et al. Prefrontal projections modulate recurrent circuitry in the insular cortex to support short-term memory[J/OL]. Cell Reports, 2024, 43(2): 113756. DOI:10.1016/j.celrep.2024.113756.

[61] English D F, Mckenzie S, Evans T, et al. Pyramidal Cell-Interneuron Circuit Architecture and Dynamics in Hippocampal Networks[J/OL]. Neuron, 2017, 96(2): 505–520.e7. DOI:10.1016/j.neuron.2017.09.033.

[62] Stark E, Abeles M. Unbiased estimation of precise temporal correlations between spike trains[J/OL]. Journal of Neuroscience Methods, 2009, 179(1): 90–100. DOI:10.1016/j.jneumeth.2008.12.029.

[63] Abeles M. Quantification, smoothing, and confidence limits for single-units’ histograms[J/OL]. Journal of Neuroscience Methods, 1982, 5(4): 317–325. DOI:10.1016/0165-0270(82)90002-4.

