## Supplementary material for "A Brain-wide Neuronal Spiking and Behavior Dataset for Working Memory-Specific Activation and Reactivation in Mice": Table S1

**Table S1. Distribution of Recorded Neurons Across Minor Brain Regions.**

| Index | Abbreviation | Name | All neurons | Memory-content-selective Neurons |
| --- | --- | --- | --- | --- |
| 35 | ATN | Anterior group of the dorsal thalamus | 86 | 18 |
| 36 | AUD | Auditory areas | 81 | 16 |
| 37 | LPO | Lateral preoptic area | 57 | 20 |
| 38 | FS | Fundus of striatum | 47 | 16 |
| 39 | RPF | Retroparafascicular nucleus | 42 | 8 |
| 40 | GU | Gustatory areas | 39 | 3 |
| 41 | NPC | Nucleus of the posterior commissure | 32 | 6 |
| 42 | AAA | Anterior amygdalar area | 29 | 11 |
| 43 | SH | Septohippocampal nucleus | 29 | 20 |
| 44 | EPv | Endopiriform nucleus ventral part | 23 | 16 |
| 45 | BMAa | Basomedial amygdalar nucleus anterior part | 22 | 5 |
| 46 | MEA | Medial amygdalar nucleus | 21 | 5 |
| 47 | IA | Intercalated amygdalar nucleus | 20 | 11 |
| 48 | PTLp | Posterior parietal association areas | 20 | 9 |
| 49 | SCig | Superior colliculus motor related intermediate gray layers | 17 | 4 |
| 50 | ECT | Ectorhinal area | 15 | 4 |
| 51 | GENv | Geniculate group ventral thalamus | 13 | 1 |
| 52 | MOB | Main olfactory bulb | 13 | 9 |
| 53 | PRC | Precommissural nucleus | 13 | 2 |
| 54 | TEa | Temporal association areas | 13 | 3 |
| 55 | PS | Parastrial nucleus | 12 | 1 |
| 56 | PP | Peripeduncular nucleus | 10 | 1 |
| 57 | OP | Olivary pretectal nucleus | 9 | 0 |
| 58 | BMAp | Basomedial amygdalar nucleus posterior part | 7 | 4 |
| 59 | MA | Magnocellular nucleus | 4 | 2 |
| 60 | NLOT | Nucleus of the lateral olfactory tract | 4 | 1 |
| 61 | BAC | Bed nucleus of the anterior commissure | 1 | 1 |
| 62 | MPO | Medial preoptic area | 1 | 0 |
